# Supervised dimensionality reduction for exploration of single-cell data by Hybrid Subset Selection - Linear Discriminant Analysis

**DOI:** 10.1101/2022.01.06.475279

**Authors:** Meelad Amouzgar, David R. Glass, Reema Baskar, Inna Averbukh, Samuel C. Kimmey, Albert G. Tsai, Felix J. Hartmann, Sean C. Bendall

## Abstract

Single-cell technologies generate large, high-dimensional datasets encompassing a diversity of omics. Dimensionality reduction enables visualization of data by representing cells in two-dimensional plots that capture the structure and heterogeneity of the original dataset. Visualizations contribute to human understanding of data and are useful for guiding both quantitative and qualitative analysis of cellular relationships. Existing algorithms are typically unsupervised, utilizing only measured features to generate manifolds, disregarding known biological labels such as cell type or experimental timepoint. Here, we repurpose the classification algorithm, linear discriminant analysis (LDA), for supervised dimensionality reduction of single-cell data. LDA identifies linear combinations of predictors that optimally separate *a priori* classes, enabling users to tailor visualizations to separate specific aspects of cellular heterogeneity. We implement feature selection by hybrid subset selection (HSS) and demonstrate that this flexible, computationally-efficient approach generates non-stochastic, interpretable axes amenable to diverse biological processes, such as differentiation over time and cell cycle. We benchmark HSS-LDA against several popular dimensionality reduction algorithms and illustrate its utility and versatility for exploration of single-cell mass cytometry, transcriptomics and chromatin accessibility data.

## Introduction

Single-cell technologies have revolutionized our understanding of biology, enabling granular dissection of the cellular heterogeneity present in complex biological samples. A surge of innovative method development has provided researchers with the means to quantify the transcriptome (Tang et al., 2009), immunophenotype (Glass et al., 2020), chromatin accessibility (Buenrostro et al., 2015), clonality (Han et al., 2014), and antigen-specificity (Newell et al., 2013) of single-cells, in some cases simultaneously (Stoeckius et al., 2017; Swanson et al., 2021). Mass cytometry (CyTOF) facilitates quantification of ~50 parameters on millions of cells in a single experiment (Bendall et al., 2011), while sequencing-based approaches can measure tens of thousands of features on tens of thousands of cells (King et al., 2021). This deluge of data encompassing a diversity of omics, cell quantities, dimensionalities, and biological samples is not amenable to a single computational pipeline or approach for analysis, but instead requires a range of flexible computational tools to address different biological questions and therefore analytical needs.

Dimensionality reduction facilitates exploration of these large, high-dimensional datasets by generating a two-dimensional coordinate system that enables simultaneous visualization of all datapoints in a single biaxial plot that captures the high-dimensional relationships of cells. Principal component analysis (PCA) performs unsupervised dimensionality reduction by identifying linear combinations of features that maximize variance (Pearson, 1901). While PCA has been applied to high-dimensional single-cell data (Newell et al., 2012), non-linear unsupervised methods, such as UMAP (Becht et al., 2019) and PHATE (Moon et al., 2019), have been widely adopted for single-cell visualization due to a superior ability to capture local and global structure, while preventing coordinate overlap. While these algorithms represent powerful tools for computational biology they may not always be the optimal choice for a given dataset, based on biological question, analysis goal, and/or available computational resources. Furthermore, while these unsupervised methods provide an unbiased view of the data, they cannot utilize *a priori* knowledge of sample composition to improve the manifold.

Previously, we introduced linear discriminant analysis (LDA) for visualization of single-cell morphometry data for hematopathology diagnostics driven by previously defined healthy cell classes (Tsai et al., 2020). LDA is a classification algorithm that identifies linear combinations of features that optimally separate previously-determined class labels (Hastie et al., 2009). LDA is primarily used to predict the class label of new observations, but we instead exploit the inherent dimensionality reduction of the method for visualization and hypothesis generation, rather than classification. Here, we demonstrate that LDA is an effective supervised tool to visualize and organize cells according to *a priori* labels such as cell type, cell cycle phase, or experimental time point. We implement hybrid subset selection (HSS), a heuristic approach utilizing elements of both forward and reverse stepwise selection, to identify a set of features that enable enhanced separation of these labels. Furthermore, feature selection by HSS for optimization of class separation combined with data visualization provides users with a visually intuitive and interpretable understanding of key feature drivers underlying the biological source of variation represented by class labels. We compare and benchmark HSS optimized LDA against PCA, UMAP, and PHATE across three mass cytometry datasets and demonstrate its utility and versatility for visualization of single-cell transcriptomics and epigenetic profiling. Finally, to empower researchers to apply supervised dimensionality reduction to their own datasets, we introduce our implementation of LDA with feature selection in the R package, *hsslda*.

## Results

### Hybrid Subset Selection optimizes supervised dimensionality reduction for single-cell visualization

Supervised dimensionality reduction by LDA takes in a matrix of cells (n) and features (p), as well as a list of *a priori* classes (k), to generate a set of k - 1 linear discriminants (LDs) (Fig. 1A). LDA leverages these class assignments as a response variable to derive the LDs, which are interpretable linear combinations of features that optimally separate cells by their known, user-defined class assignment. These *a priori* labels can be biological features of cells such as cell-types, collection timepoints, cell lines, cell cycle phases, or other categorical/ordinal features. Traditionally, dimensionality reduction relies on all defined features (p) as inputs. However, to obtain the optimal separation between classes for visualization, the user needs to tune this feature set so that it best separates the class labels in the data. This separate analysis can often be a time-intensive task for biologists. To facilitate improved dimensionality reduction and visualization, we implemented HSS to augment LDA with an automatic feature selection that optimizes class separation in an interpretable manner (Fig. 1B). The HSS-LDA algorithm uses a combination forward and reverse stepwise feature selection heuristic, calculating separation scores across many feature subsets, and selects the final set of features that best separates classes for visualization (see *Methods*).

**Figure 1:**
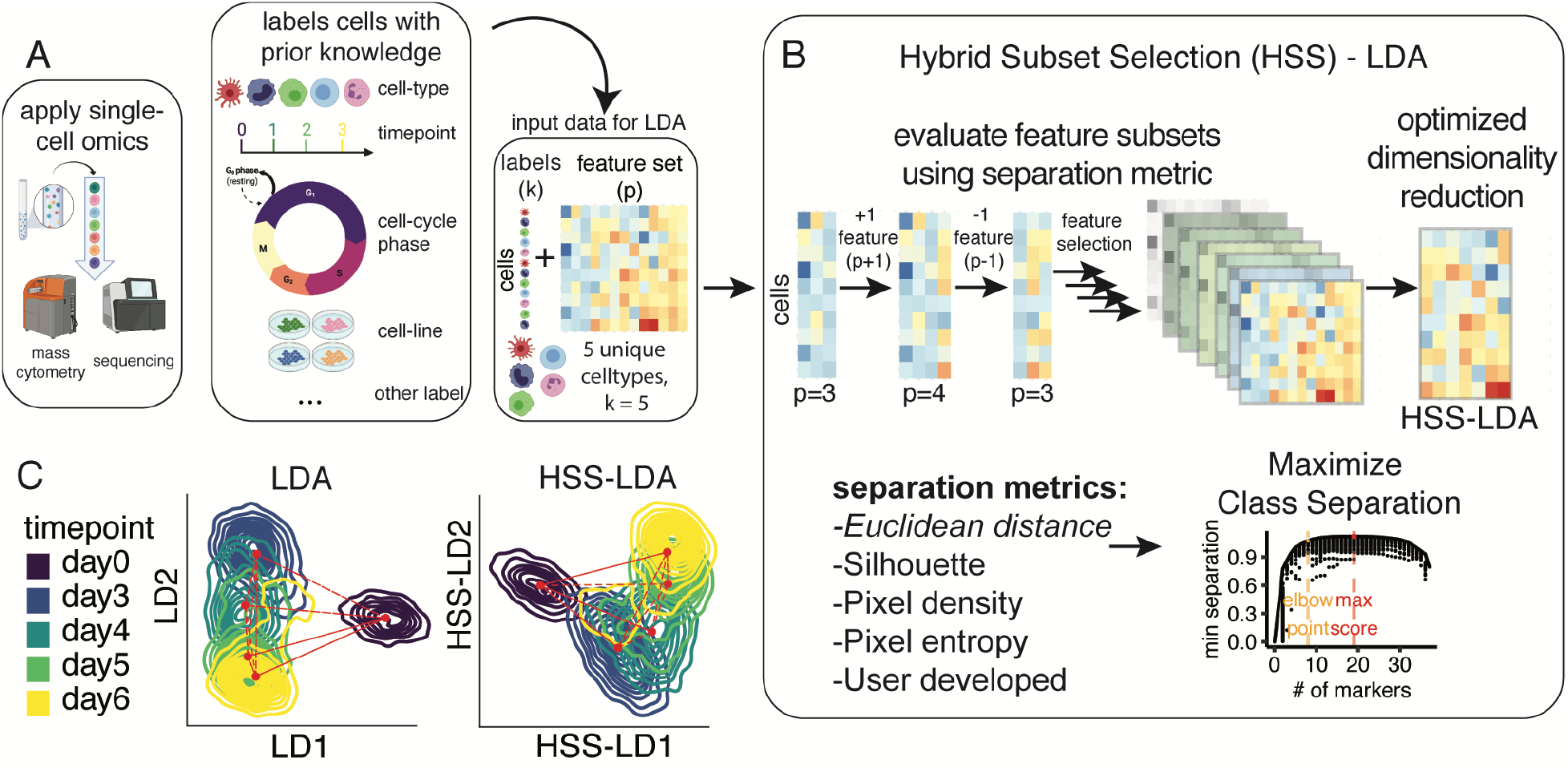
HSS-LDA optimizes dimensionality reduction using feature selection. **(A)** Workflow demonstrating Linear Discriminant Analysis (LDA) with prior knowledge of class labels of interest for supervised dimensionality reduction and feature selection using Hybrid-Subset-Selection (HSS). **(B)** HSS-LDA performs feature selection to enhance dimensionality reduction and visualization of single-cell data by maximizing class separation via a stepwise feature selection approach, selecting the final model based on a separation metric specified by the user. **(C)** Comparison of LDA and HSS-LDA visualization using example endoderm differentiation data.

For example, to understand endoderm differentiation patterns, Kimmey *et al*. collected endoderm cells across five differentiation timepoints (k=5) and applied CyTOF to obtain a single-cell matrix of cells and protein markers (Kimmey et al., 2022 - BioRxiv in process). In addition to this single-cell matrix of protein features, each cell was also annotated with its collection timepoint. By applying LDA and visualizing the first two LDs, cells from different timepoints were separated across a single biaxial plot (Fig. 1C, left). Day 0 separated from the other time points but its relationship to other timepoints in differentiation was not evident. While LDA alone can visually separate cells according to their class assignments, we applied HSS to identify the combination of features that optimizes this separation, resulting in improved visualization (Fig. 1C, right). HSS-LDA improved differentiation timepoints with an optimized set of features and revealed a continuous trajectory from hESC day 0 to day 6 differentiated endoderm-primed cells.

To assess the utility of HSS-LDA for single-cell visualization, we applied LDA, HSS-LDA, and UMAP (Becht et al., 2019) to three CyTOF datasets from different biological systems and with different visualization needs. Each dataset was assigned a specific name for this paper: “Morphometry” (Tsai et al., 2020), “T-cell metabolic regulome” (Hartmann et al., 2020), and “Chromotyping” (Baskar et al., BioRxiv in process). These mass cytometry datasets had unique challenges such as significantly imbalanced cell numbers across classes (eg, celltypes), as well as different biological processes to visualize such as discrete, continuous, or cyclical systems. Compared to LDA, the HSS-LDA visualization either qualitatively improved or retained the original separation of class labels in the HSS-LDA embedding for the three datasets despite using a lower dimensional feature matrix (Fig. S1). HSS-LDA facilitated separation of cell-types in the morphometry dataset using only 13 features from 17 (Fig. S1A-B), metabolic states of T-cells over time using only 8 features from 32 (Fig. S1C-D), and cell cycle phases using only 24 features from 32 (Fig. S1E-F). In addition to identifying an optimized minimum feature set through feature selection to reduce the feature space, the HSS-LD coefficients also provide interpretability of key drivers of class separation in both magnitude and direction, which can be used to guide other analyses.

### HSS-LDA reconstructs both discrete and continuous biological processes

The Morphometry dataset employs a set of markers coined ‘scatterbodies’ that capture immune cellular identities based on structural features that are consistent even in malignancy thereby discriminating immune and hematopoietic cells extracted from bone marrow (Fig. 2A). There are often drastic cell type imbalances in human tissues, exemplified in the Morphometry dataset by a predominance of neutrophils compared to other cell populations in the bone marrow (Fig. 2B-C). While this can be re-balanced through equal sampling of each cell-type, subsetting the data potentially discards valuable cell information reliant on prior system knowledge. UMAP spreads data to avoid coordinate overlap, but in this imbalanced dataset, the result is neutrophils dominating the entire manifold, making it difficult to see differences between classes. HSS-LDA treats discrete cell-types equally and separated cell-types regardless of their cell abundance, while preserving cellular relationships by protein abundance (Fig. 2C, S2A-B). To test the HSS-LDA’s function as a classification algorithm, we trained HSS-LDA for both visualization and classification of discrete cellular identities using only cells from healthy donors (Fig. S3A). We visualized HSS-LDA plots and accuracy metrics, finding that HSS-LDA accurately predicted cellular identities of cells derived from patients with hematopoietic malignancies, with a median accuracy of ~90% (Fig. S3B-C). Thus, both the visualization and classification aspects of HSS-LDA bear utility for biological applications.

**Figure 2:**
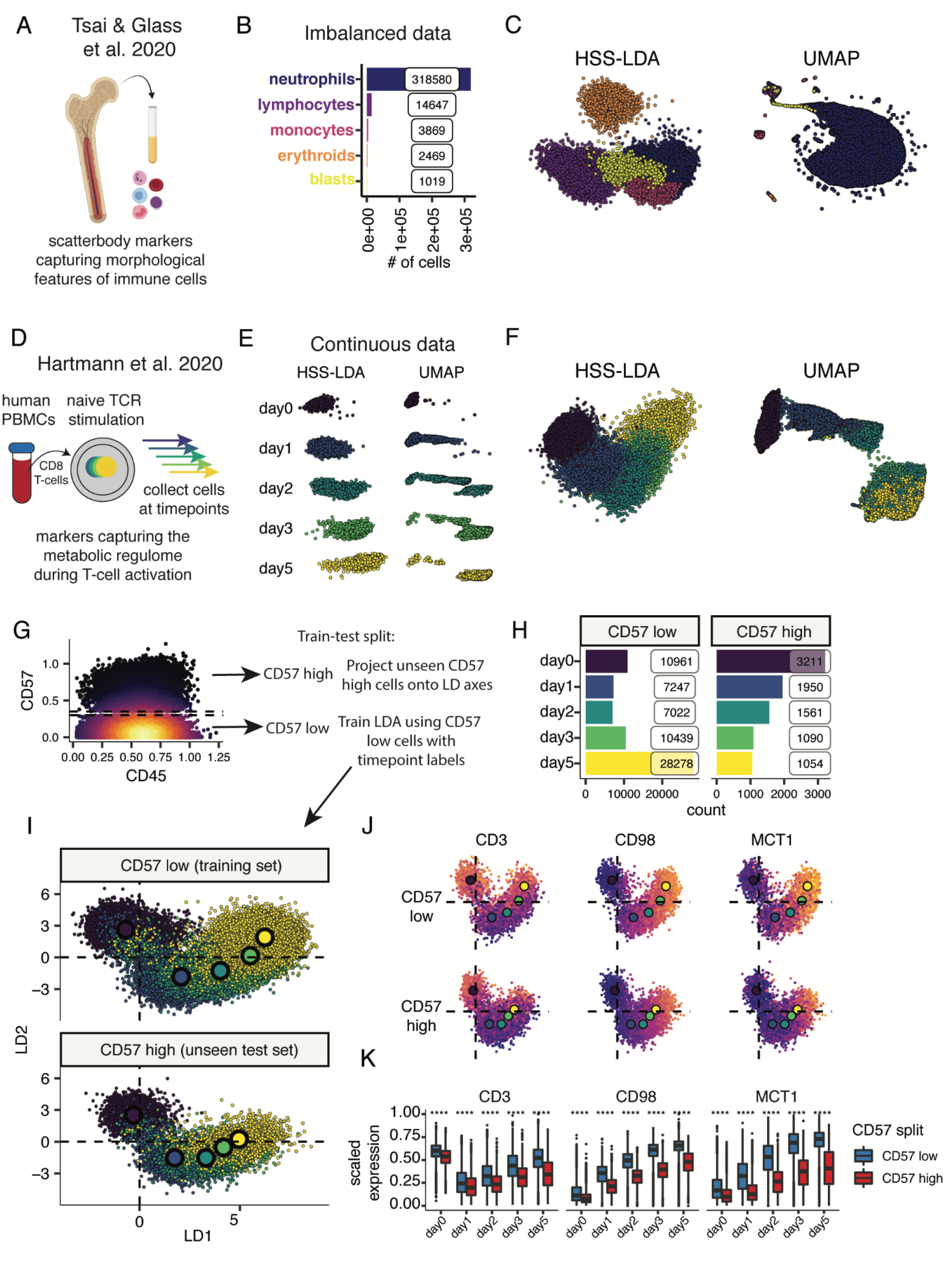
HSS-LDA reconstructs both discrete and continuous biological processes, and can embed new, unseen cells onto the visualization for exploratory analysis. **(A)** Conceptual diagram of immune cells extracted from healthy bone marrow and stained using morphometric markers for CyTOF. **(B)** Barplot summarizing imbalanced class distribution of immune cell populations. **(C)** Comparison of HSS-LDA using PCE score for feature selection and UMAP demonstrating discrete class visualization using the same input cells. **(D)** Conceptual diagram of human CD8 naive T cells extracted from PBMCs for *in vitro* TCR stimulation, collected on days 0-5 of activation, and stained with metabolic markers for CyTOF analysis. **(E)** Comparison of HSS-LDA using euclidean distance for feature selection and UMAP demonstrating a linear trajectory using the same input cells faceted across each timepoint. The # of cells are balanced across each timepoint. UMAP implemented with published settings: n_neighbors = 15 and min_dist = 0.02. **(F)** Unfacetted HSS-LDA and UMAP plot of figure 2*E*. **(G)** Biaxial CD57 versus CD45 plot showing train-test split for stratification CD57^low^ and CD57^high^ cells. HSS-LDA is trained on CD57^low^ cells and the unseen CD57^high^ cells are used as a test set projected onto the CD57^low^ LD embedding. **(H)** Barplot summary counts for CD57^low^ and CD57^high^ training and test sets. **(I)** Biaxial LD plots of CD57^low^ cells and embedded CD57^high^ cells labeled with the centroid point for each timepoint. **(J)** Protein expression of biaxial HSS-LD plots for three example markers: CD3, CD98, and MCT1. **(K)** Boxplot summary of protein expression for CD57^low^ and CD57^high^ cells across each timepoint. Wilcoxon signed-rank test performed between CD57^low^ and CD57^high^ cells across each timepoint. *: p <= 0.05; **: p <= 0.01; ***: p <= 0.001; ****: p <= 0.0001.

The T-cell metabolic regulome dataset includes a feature set of markers that capture the metabolic state of human peripheral blood mononuclear cell (PBMC)-derived CD8+ T-cells collected across multiple timepoints after *in vitro* TCR stimulation (Fig. 2D). Hartmann *et al*. demonstrated that markers for metabolic regulation capture the continuous trajectory of metabolic cellular states over time. Both HSS-LDA and UMAP separated cells across each timepoint and visualized the linear trajectory of T-cell metabolism after TCR stimulation (Fig. 2E-F, Fig. S2C-D). LDA also provided both (1) interpretable feature coefficients across each linear discriminant, and (2) facilitated projection of unseen data onto previously trained LD axes for exploratory analysis. We demonstrated this utility by stratifying T-cells by CD57 expression - a marker of senescence and terminal differentiation (Fig. 2G-H). We trained LDA on CD57^low^ cells and visualized both the linear trajectory of these cells and the respective linear combination of coefficients in a biaxial plot to generate a latent space representation of the metabolic progression of a non-senescent T cell during TCR stimulation (Fig. 2I). The largest contributors to LD1 and LD2 that separate stimulated T-cells over time were CD98, OGDH, GLUD1/2, CS, MCT1, and GLUT1 (Fig. S3D-F). To visualize the metabolic progression of senescent T-cells compared to non-senescent T cells during TCR stimulation, we projected the unseen CD57^high^ cells onto the CD57^low^ LD axes and found that CD57^high^ cells had a stunted metabolic progression starting between days 1-2 of TCR stimulation compared to CD57^low^ cells (Fig. 2I). The mean coordinates of CD57^high^ cells at day 5 overlapped on the manifold with the CD57^low^ cells at around days 2-3 of TCR stimulation, reflecting that these cells shared a common metabolic state on different days of TCR stimulation, reflected by their metabolic protein abundance (Fig. 2J-K). We explicitly assessed all metabolic markers and implemented LDA rather than HSS-LDA to train a more metabolically integrative model that is less biased towards a CD57^low^-specific feature set. Through supervised dimensionality reduction by LDA, we show that a supervised method can be trained on a baseline cellular state such as T-cells with healthy proliferation potential, and distinct cellular states such as those with senescent or terminally differentiated phenotypes can be compared to a baseline state while integrating high-dimensional data in a visually intuitive manner. These results demonstrate that LDA and HSS-LDA can be applied to datasets with both discrete categorical and continuous trajectory labels with overlapping features to assist in visualizing and interpreting single-cell biological systems.

### HSS-LDA captures cyclical biological processes within multi-label data

Cellular division is an important and highly regulated biological process which maintains tissue homeostasis with cellular turnover and can become corrupted in malignancy. Through cellular division, global chromatin structure undergoes significant changes to facilitate DNA replication and separation into 2 cells. To better understand the dynamics of chromatin structure regulators through the cell cycle, we applied HSS-LDA to highly multiplexed chromatin content data from single cells (i.e. chromotype: a collection of chromatin-modifying factors and histone modifications) across cell lines and cell cycle states (Fig. 3A-B). Global chromatin content as defined by single cell abundance of chromatin-modifying factors and histone modifications capture the distinct, endogenous epigenetic patterning of different cell lines as well as the expression patterns of these markers across the cell cycle (Fig. 3A). This dataset is particularly unique because it contains (1) two sets of labels (cell-type and cell cycle), and (2) a cyclical biological process (cell cycle), resulting in five cell lines and five cell cycle phases (Fig. 3B). Both the cell lines and cell cycle phases contain an imbalanced distribution of cells. Ground truth cell cycle phases were labeled by manual gating of CyclinB1, IdU, phosphorylated H3, and pRB (Kimmey et al., 2019, Bartek et al., 1997).

**Figure 3:**
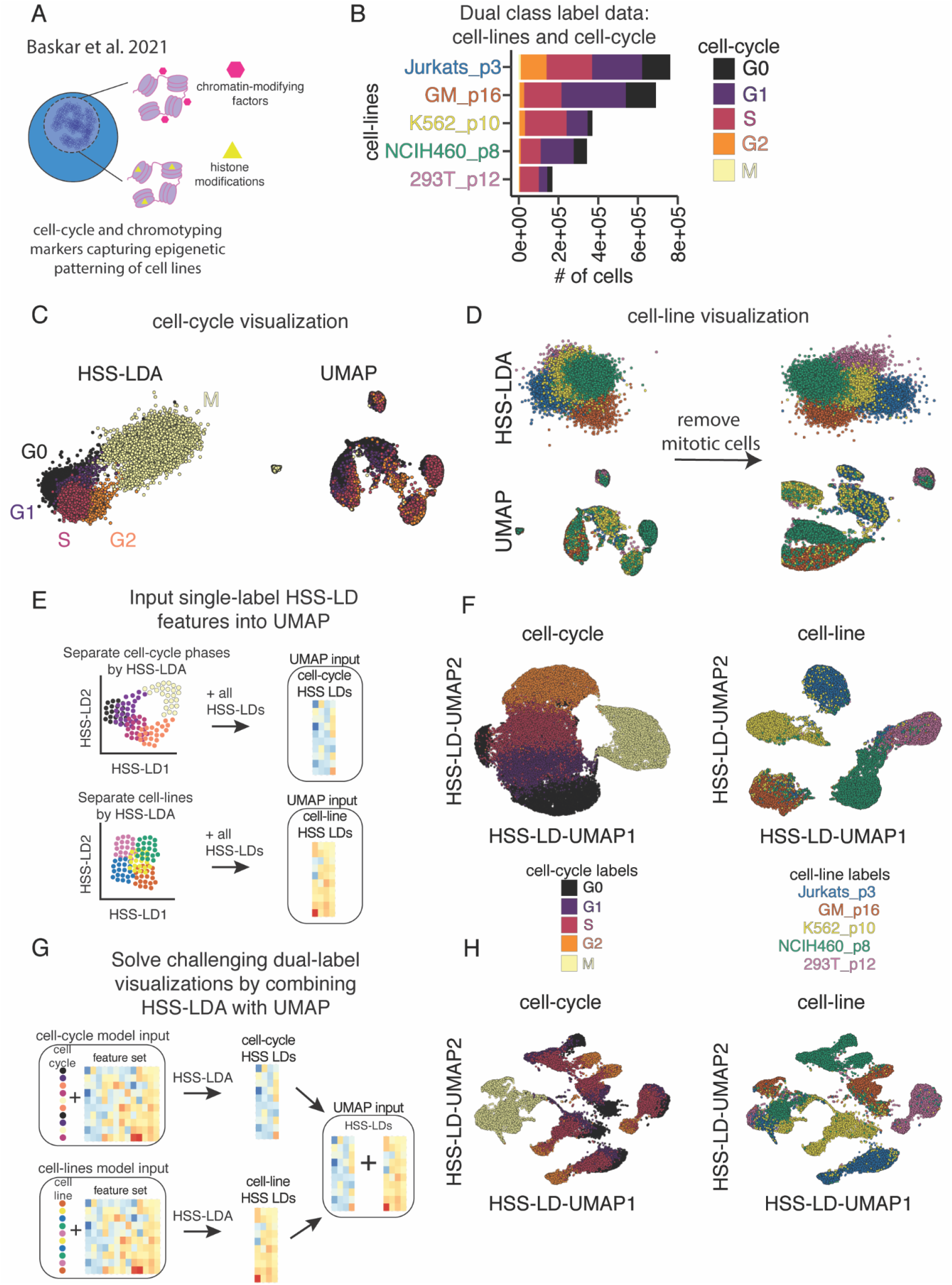
HSS-LDA reconstructs cyclical biological trajectories and can be input as features into UMAP to solve challenging dual-class visualization tasks. **(A)** Conceptual diagram of cell cycle and chromotyping markers of various cell lines for CyTOF analysis. **(B)** Barplot summary of cell counts for each cell line in various cell cycle phases. **(C)** Comparison of HSS-LDA using euclidean distance for feature selection and UMAP visualizing the cell cycle. **(D)** Comparison of HSS-LDA using euclidean distance for feature selection and UMAP both including and excluding mitotic cells to visualize cell lines. **(E)** Conceptual diagram demonstrating prior supervised dimensionality reduction using HSS-LDA to initialize UMAP. **(F)** HSS-LDA-initialized UMAP plots of the cell cycle and cell line labels. UMAP parameters for cell cycle are n_neighbors = 25, spread= 7, and cell lines are n_neighbors = 15, spread= 1. **(G)** Conceptual diagram demonstrating prior supervised dimensionality reduction using HSS-LDA to initialize UMAP for dual-class labelled data visualization. HSS-LDA is computed separately on cell cycle and cell line labels, and the HSS-LDs are merged as the feature set input to initialize UMAP. **(H)** HSS-LDA-initialized UMAP plots demonstrating dual-class visualization of both cell line and cell cycle systems in a single biaxial plot. UMAP parameters are n_neighbors = 10, spread= 4.

Cell cycle is thought of as a circular process where the M-phase parental cell divides to form two G0/G1 child cells. Here, we wanted to explicitly visualize this cyclical trajectory to track protein expression across the cell cycle independent of cell line heterogeneity. We ran HSS-LDA and UMAP using the same initial set of epigenetic markers and trained HSS-LDA using the cell cycle labels. HSS-LDA successfully separated the cell cycle states as well as captured the circular trajectory of the cell cycle, whereas UMAP did not separate cell cycle labels as an adequate representation of the cell cycle stages (Fig. 3C, S2E-F). The UMAP manifold was confounded by the imbalanced distribution of the dataset and attempted to capture information unique to both the cell lines and cell cycle. HSS-LDA selected features that separate cell cycle labels and generated an interpretable linear combination of features that separates these cell labels (Fig. S2E-F). UMAP would require manual intervention to identify the feature subset to adequately visualize the cell cycle, and even then, the cell cycle signal may still be confounded by distinct cell line properties that may require signal correction methods to deconvolve cell cycle and cell line properties. The mitotic cells were hidden in the UMAP and other cell cycle phases were projected onto multiple areas of the manifold. While parameter tuning could perhaps improve the UMAP embedding, it cannot fully resolve these visualization challenges. HSS-LDA takes advantage of prior knowledge of the cell cycle labels and feature selection captures the relevant cell cycle information independent of the cell lines, sufficiently visualizing the cyclical trajectory using a linear transformation. Here, we can visibly see marker transitions of puromycin and CyclinB1 protein abundances along the cell cycle trajectory (Fig. S2E). By visualizing cellular relationships in a manner that reflects the cyclical trajectory of the cell cycle, we can more intuitively study the markers that directly contribute to cell cycle dynamics but also independently study the patterns of other protein markers that are not used to construct the cell cycle embedding.

At the same time, within the same dataset HSS-LDA could reveal cell line differences, visualizing them independent of cell cycle phases. While projecting the same data using cell line labels proves more difficult for both HSS-LDA and UMAP, the HSS-LDA plot further improved when mitotic cells were removed (Fig. 3D, Fig. S2G-H). However, this remains a challenging biaxial visualization task for both HSS-LDA and UMAP, which is one of the current gold-standard dimensionality reduction methods for visualization. For both cell cycle and cell line, low-abundance classes (eg, mitotic cells) were better visualized with HSS-LDA as compared to UMAP (Fig. 3C-D), as they occupy a proportionally larger area on the manifold in UMAP, as previously shown with the Morphometry dataset. Thus, HSS-LDA can be used to visualize cyclical biological trajectories and discrete cellular identities with heterogeneous data distributions, enabling identification of features uniquely associated with either cell cycle or cell line classes.

### HSS-LDA as UMAP input integrates variance from multiple class labels into a single visualization

While we utilized HSS-LDA as a visualization tool using the first two Linear Discriminants (LDs), HSS-LDA produces multiple LDs. The number of LDs produced in the model is calculated as the [# of classes - 1]. Subsequent HSS-LDs contain additional information that separate classes of interest. To capture data patterns resulting from more than one source of known variance, we hypothesized that HSS-LDs generated from multiple class labels could be used as input to unsupervised dimensionality reduction methods. We performed supervised dimensionality reduction by HSS-LDA for either cell cycle or cell line labels and inputted the HSS-LDs as features into UMAP to generate an HSS-LD-UMAP embedding that sufficiently separates classes (Fig. 3E-F).

Given that HSS-LD-UMAP embeddings can separate classes within each single label, we tested if we could exploit a combination approach to dimensionality reduction to visualize both cell line and cell cycle class labels in a single biaxial plot. We combined the HSS-LDs from the two separate HSS-LDA analysis for cell line and cell cycle into a single table and inputted this as a feature set into UMAP to generate a combinatorial HSS-LD-UMAP embedding (Fig. 3G). The resulting biaxial plot preserves both cell line and cell cycle relationships in a biologically meaningful manner (Fig. 3H). Cell lines cluster separately (Fig. 3H, *right panel*) while still preserving the cell cycle trajectory from the G0-G2 state within each cell line (Fig. 3H, left panel).

Cell line differences are more distinct than cell cycle differences, with major patterns being driven by basal epigenetic differences between cell lines. Expectedly, unlike all other cell cycle states, global chromatin content of cells in mitotic phase is highly conserved across cell lines. As shown, dual-label visualizations can be useful to demonstrate distinct patterns in a biological process across different systems. Apart from better representing multiple known sources of heterogeneity in a single embedding, prior supervised dimensionality reduction by LDA also significantly reduces the number of features inputted into UMAP from a full panel of markers to a smaller set of HSS-LDs, reducing UMAP runtime on large single cell datasets. This is similar to one of many advantages when performing PCA to reduce the feature set prior to UMAP in high-dimensional genomics datasets. Thus, HSS-LDA for supervised dimensionality reduction can be used prior to unsupervised methods like UMAP to help solve challenging dual-label and multi-class visualization tasks in a single embedding.

### Benchmarking LDA efficiency and class separation against common dimensionality reduction algorithms

To further assess the utility of LDA and HSS-LDA, we extended our comparison to other popular dimensionality reduction methods, including PCA (Pearson, 1901), UMAP (Becht et al., 2019), and PHATE (Moon et al., 2019) across the three mass cytometry datasets (Fig. 4A). PCA is an ideal comparison because it is conceptually similar to LDA in that they both are linear transformation techniques. While PCA finds directions of maximal variance, LDA finds the feature subspace that maximizes class separability. UMAP is a nonlinear, dimensionality-reduction technique that is arguably the most popular single-cell visualization tool, and computationally shares many similarities with its related predecessor tSNE (Kobak & Linderman, 2021). PHATE, more recently introduced, is an information-geometric distance approach to capture local and global nonlinear structure for dimensionality reduction.

**Figure 4:**
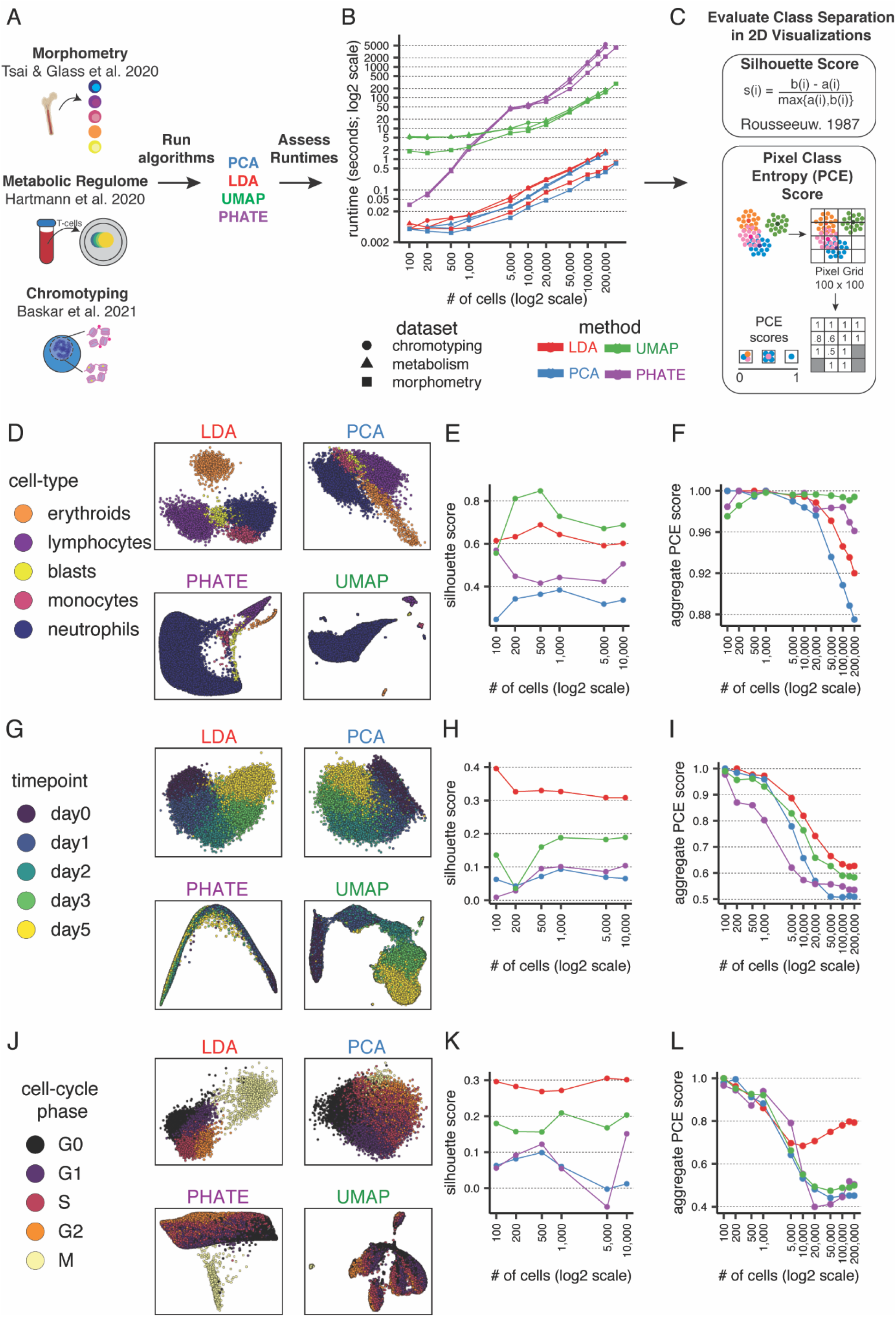
LDA is computationally efficient, scalable, and adequately separates class labels. **(A)** Conceptual diagram for comparing various dimensionality reduction algorithms. PCA, LDA, UMAP, and PHATE algorithms are applied to three CyTOF datasets and runtimes are assessed to determine efficiency and scalability of the algorithm. **(B)** The average runtime of three analyses across three datasets for each algorithm are shown across different dataset sizes on a log2-transformed scale. Default algorithm settings are used. **(C)** Summary of silhouette score and Pixel Class Entropy (PCE) score to assess separation of class labels of interest for each algorithm. Both metrics can be used for feature selection by HSS-LDA. **(D-L)** Summary plots of each algorithm applied to the morphometry, T-cell metabolism, and chromotyping datasets. (*left*: *D, G, J*) Representative biaxial visualizations of each algorithm using 10,000 cells. (*middle*: *E, H, K*) Average silhouette score across different cell counts for each algorithm. (right: *F, I, L*) Average PCE score in a 100 x 100 pixel grid across different cell counts for each algorithm.

We compared LDA to these diverse algorithms to emphasize use-cases where one algorithm may be more suitable than another. To fairly compare runtimes, we used the same feature set and input cells for each algorithm. As anticipated, LDA and PCA were significantly faster algorithms than UMAP and PHATE. We emphasize that a log2 scale was required to adequately visualize the runtime discrepancy between the algorithms as cell counts increased (Fig. 4B). LDA is non-stochastic, reproducible, and robust with even low cell counts. Visualizing all four algorithms across the varying cell counts demonstrated that cells will generally occupy the same phenotypic coordinates in the LD embedding when using LDA (Fig. S4A-C). LDA’s scalability makes it amenable for feature selection using our hybrid subset selection approach. HSS-LDA runtimes converge with other dimensionality reduction methods that do not provide feature selection when reaching a range of cells that can accurately identify a minimum feature set (Fig. S5A-C). Using the three mass cytometry datasets, we identified a heuristic of approximately 100-200k cells to be used as a subset prior to running HSS-LDA to identify the optimal feature set, though there will be variability with every dataset (Fig. S5D). Once the HSS-LDs are computed, the model can non-stochastically project the same data to reproduce the exact same axes or project unseen data within seconds, as was done in the T-cell metabolic regulome dataset training the LDA model on the CD57^low^ cells and projecting the CD57^high^ cells onto the same LD axes for rapid dimensionality reduction (Fig. 3G-K).

To quantitatively assess class label separation, we varied cell count inputs into each algorithm and applied two separation metrics: (1) silhouette score and (2) pixel class entropy (PCE) score (Fig. 4C). Silhouette score is a measure of how similar cells are to their own cluster compared to other clusters by accounting for both intra-cluster and inter-cluster Euclidean distance of each class. PCE score pixelates the biaxial plot into a grid and computes an average PCE score measured by the entropy of all class labels in each pixel of the grid - the approach is further described in the methods. To fairly assess the four algorithms, we used the final feature set selected using HSS-LDA as input into PCA, UMAP, PHATE, and LDA.

A 50k cell subset of the three mass cytometry datasets and their respective labels (Fig.4D, G, J - *left*) were embedded using these methods. We found that LDA adequately separated class labels across the three datasets and often performed better compared to other dimensionality reduction algorithms. Silhouette score summaries related that LDA performed second best to UMAP in separating cell-types in the Morphometry dataset, though as demonstrated in Fig. 2, UMAP failed to handle data imbalances and yielded a less useful visualization due to underrepresentation of rare cell types. LDA performed the best separating intra-cluster and inter-cluster distance across timepoints in the T-cell metabolic regulome, and cell cycle phases in the Chromotyping datasets compared to the other algorithms (Fig. 4E, H, K - *center*).

When comparing PCE scores in Morphometry, UMAP and PHATE performed better than LDA, particularly at the highest cell inputs (Fig. 4F, I, L - *right*). However, in the less discretized, more continuous T cell activation and cell cycle datasets, PCE scores for LDA were superior. We concluded that LDA is suitable for visualizing diverse biological systems, is robust to data imbalances, and may be a preferred dimensionality reduction algorithm depending on the visualization needs when *a priori* knowledge of a class label is known – particularly in ordered and continuous datasets.

### LDA captures the trajectory of enterocyte differentiation by single-cell transcriptomics

Given the particular performance of LDA and HSS-LDA in summarizing ordered progressions in single-cell mass cytometry data, we asked whether it could have utility for a similar process mapped by single-cell RNA-sequencing (scRNAseq) as well. We applied supervised dimensionality reduction by LDA to a spatially reconstructed scRNAseq dataset of enterocytes of the intestinal villi (Fig. 5A). Moor *et al*. used spatial transcriptomics to identify gene sets that corresponded to the differentiation patterns of enterocytes across the intestinal villi, then used these validated gene sets to generate a spatially reconstructed scRNAseq dataset of enterocytes with zone labels that correspond to their location and differentiation state (Moor et al., 2018). To test whether LDA could reconstruct the linear trajectory of enterocyte differentiation, we took the prior zone labels and the first 50 principal components as the input feature set into LDA. LDA improved the linear trajectory visualization of cells in a biologically relevant manner when compared to UMAP (Fig. 5B-C). We found that expression of landmark genes with distinct spatial patterns across the intestinal villi were reflected in the arrangement of cells in LDA (Fig. 5B-C). ADA at the villus tip and SLC2A2 mid-villus demonstrated biologically relevant expression patterns that were less coherent in the UMAP, but were preserved with LDA. These findings demonstrate the utility of LDA for visualization of scRNAseq data.

**Figure 5:**
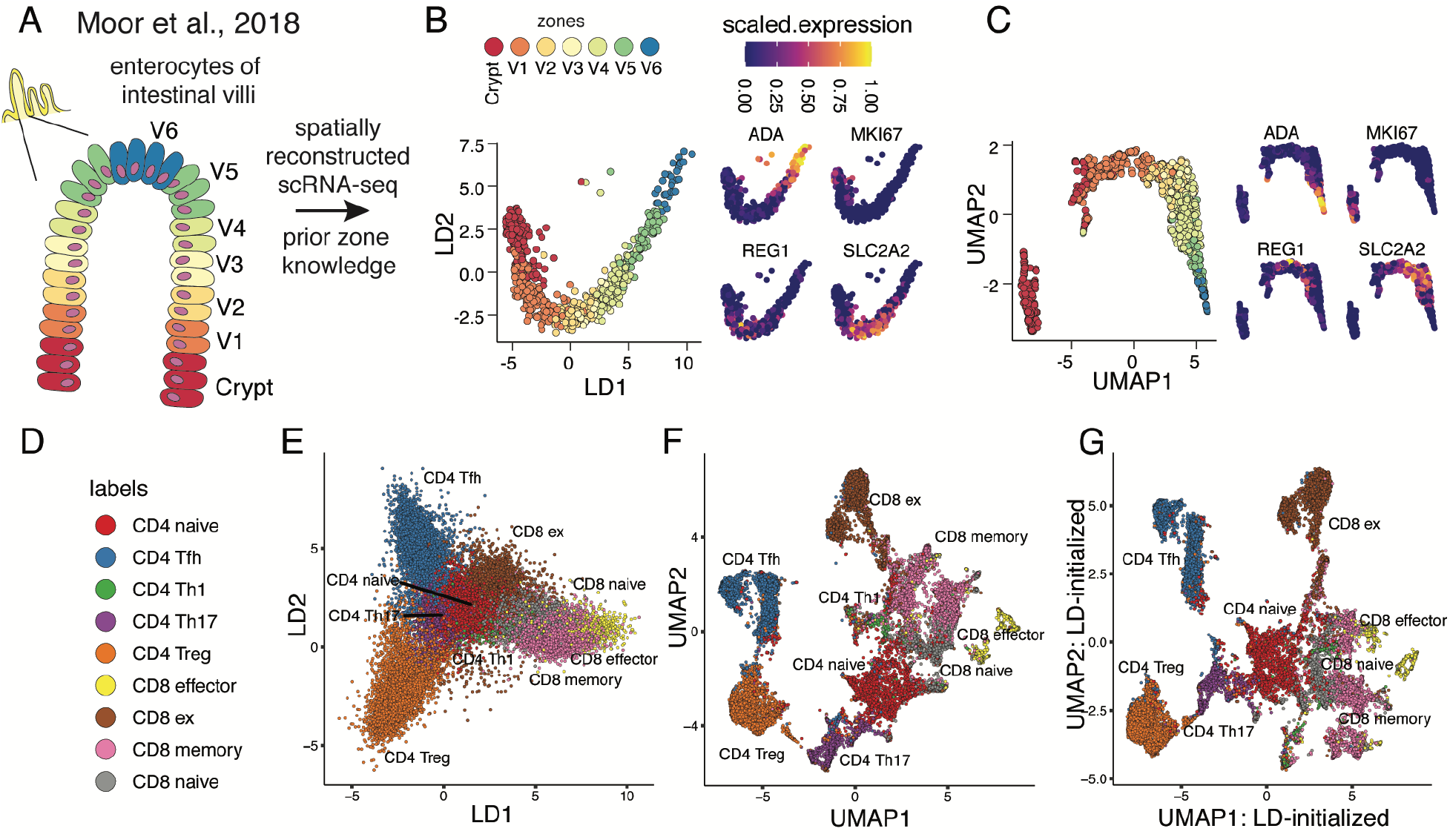
LDA utility extends to single-cell sequencing data to reconstruct linear trajectories as well as organize single-cell chromatin accessibility data using semi-supervised dimensionality reduction. **(A)** Conceptual diagram for (*B, C*) showing enterocyte differentiation from the crypt and across the intestinal villi with prior intestinal zones identified using spatial transcriptomics. **(B, C)** Comparison of LDA and UMAP demonstrating the linear trajectory of enterocyte differentiation paired with scaled expression of key genes. **(D)** Cell type labels for single-cell ATAC chromatin accessibility data of T cells from Satpathy et al., 2019 with prior known cell type labels from sorting. **(E)** LDA embedding supervised with prior known cell type labels. **(F)** UMAP embedding of the same feature set inputted in (*E*). **(G)** UMAP embedding initialized by the first 2 LDs (from *E*) using the *init* function in uwot::umap in R for semi-supervised dimensionality reduction.

### LDA organizes T cell heterogeneity seen by single-cell chromatin accessibility analysis

To extend the diversity of single-cell sequencing data types LDA could be applied to, we also investigated its utility on a single-cell assay for transposase-accessible chromatin with sequencing (scATACseq) dataset of CD4 and C8 T populations (Satpathy et al., 2019). Here, T cell populations include naive, memory, helper, effector, and exhausted subsets as sorted and annotated by the original authors. We processed ATAC peaks using the same methods as the original authors. We performed LDA supervised with celltype labels, and UMAP using the same input matrix. Both LDA and UMAP separated cell-type labels in a biologically meaningful manner (Fig. 5D-F). Inactive cell-states, such as CD4 and CD8 naive T-cells, clustered together while effector cell-types were more distant in phenotypic space. While UMAP effectively separated CD4 and CD8 subtypes and preserved nuanced single-cell relationships, LDA better preserved global structure, illustrating how these cells are related and interconnected to one another. For example, CD8 exhausted cells are embedded near the CD4 and CD8 naive cells but remain distant from CD8 effector cells. This corroborates with the more quiescent-like phenotypes of CD8 exhausted cells relative to CD8 effector cells. Kobak and Linerman previously demonstrated the importance of non-random initialization with UMAP and tSNE for generating more meaningful embeddings (Kobak & Linderman, 2021). To better preserve global relationships while exploiting the local separation in the UMAP embedding, we leverage celltype labels using LDA and initialized UMAP with LD1 and LD2 coordinates. The LD-initialized UMAP better preserved the global relationships of both the progression to CD8 exhausted states as well as the inter-relation between the different naive cell fractions compared to the un-initialized UMAP (Fig. 5G). Furthermore, we see separation of CD8 memory cells into three states in UMAP-LDA embedding compared to UMAP which underlines the utility of our tool in more than separating known class labels. Thus, LDA for supervised dimensionality reduction, can be used prior to unsupervised methods like UMAP for semi-supervised dimensionality reduction by LD-initialization to provide more useful latent space representations. Additionally, these findings demonstrate the broad utility of single cell data types LDA can robustly visualize and summarize, including scATACseq data.

### Reconstructing scRNAseq-based cell cycle pseudotime of activated human T cells using LDA

Given LDA’s utility to capture trajectories in continuous datasets (Fig. 2D-K, 5A-C) and act as in input for other embedding methods (Fig. 3), we tested whether it could help visualize a continuous, circular biological process where the classes were based on a score derived for single cell sequencing information. We curated an *ex vivo* TCR stimulation dataset that we call “T-cell Proliferation Tracing” (Good et al., 2019). This dataset contains primary human T cells labeled with CFSE, stimulated for three days *in vitro*, and prospectively isolated based on cell division state (i.e. 0 divisions, 1 division, 2 divisions) for scRNAseq analysis.

To summarize the entire cell cycle process using LDA, we computed the cell cycle phase scores for G1.S, S, G2, G2.M, and M.G1 using previously published methods (Macosko et al., 2015; Schwabe et al., 2020; Whitfield et al., 2002) (Fig. 6A). Cell cycle scores were calculated using a curated list of genes with known increased enrichment in each phase. A cell was given a score for each phase of the cell cycle, and the phase with the largest score was the cell’s assigned cell cycle state (Fig. 6B-C). However, these cell cycle phases are not entirely discrete processes from one another, which was reflected by the correlation seen between adjacent phases, driven by cells transitioning between phases (Fig. 6D). At the same time, non-adjacent phases, which had no cells transitioning between them, were anti-correlated. We applied LDA to a matrix of cell cycle scores and provided the list of cell cycle labels, resulting in a cyclical LDA visualization that accurately separated the cell cycle phases according to their expected position (Fig. 6E).

**Figure 6:**
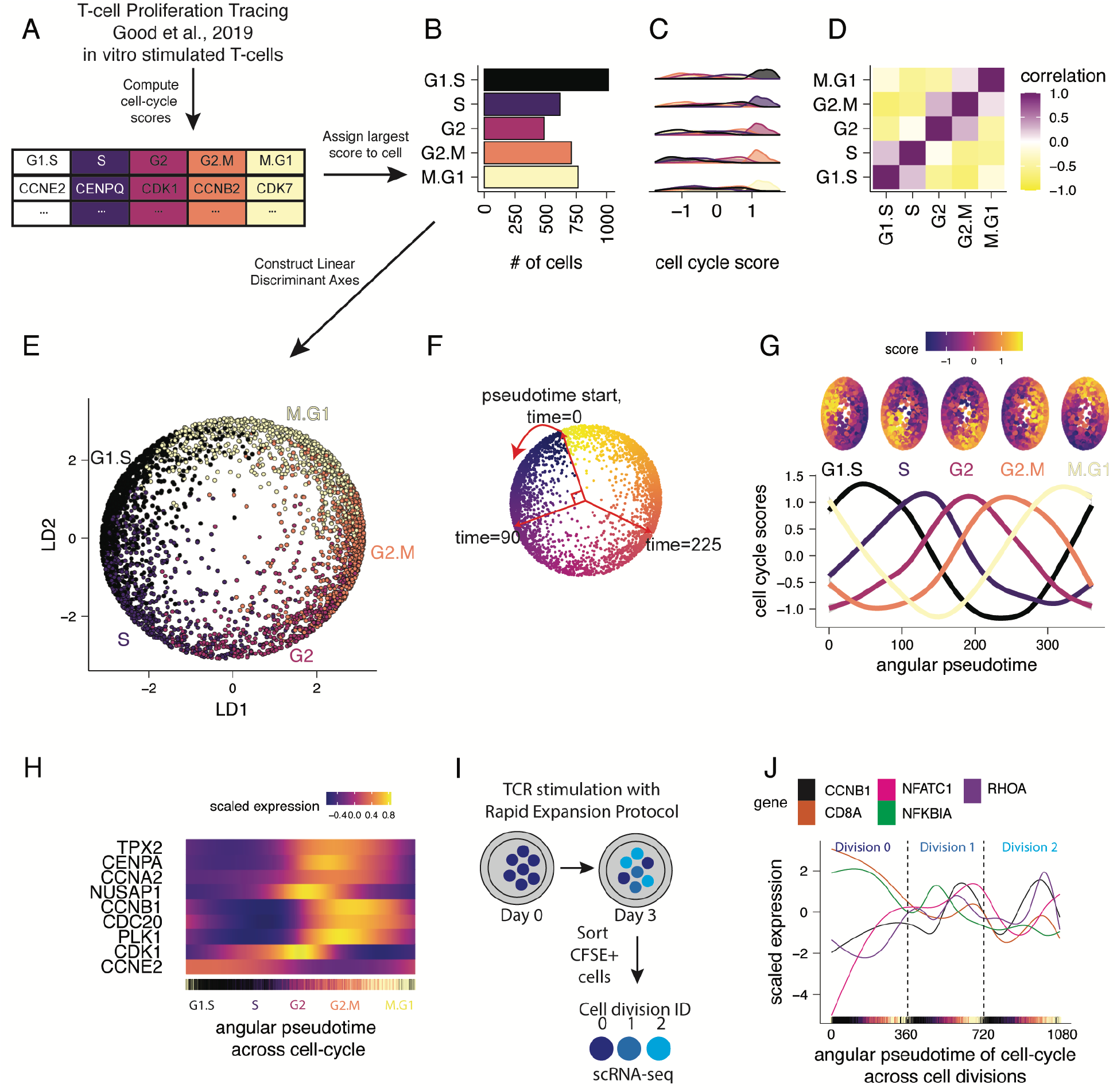
LDA can reconstruct cyclical trajectories using single-cell RNAseq data. **(A)** Conceptual diagram of cell cycle score computation using prior methods on *in vitro* CD8 T-cell TCR stimulation sc-RNAseq data. **(B)** Barplot summary counts of assigned cell cycle phase identities. The phase with the largest cell cycle score is assigned to each cell. **(C)** Density plot summary of cell cycle phases showing enrichment of cell cycle scores in each respective assigned phase. **(D)** Pearson correlation of cell cycle scores computed across all cells. **(E)** Cyclical LDA visualization on cell cycle scores. **(F)** Graphical representation of angular pseudotime calculation. **(G)** Generalized linear models of cell cycle scores across the angular pseudotime estimated using the LD biaxial. **(H)** Heatmap of estimated transcript expression summarized as a generalized additive model for key cell cycle markers across the cell cycle angular pseudotime. **(I)** Conceptual diagram for (*L*) demonstrating experimental protocol using CFSE-sorted T-cells on day 3 of TCR stimulation to extract cell division IDs prior to 10x Genomics scRNAseq. **(J)** Estimated transcript expression of CyclinB1 and relevant TCR signaling genes identified using derivative analysis plotted across the cell cycle angular pseudotime deconvolved across cell divisions using CFSE-sorted division IDs.

To assess whether the model was overfitting to the cell cycle scores, we performed cross-validation by splitting the dataset into 10 non-overlapping test sets, training the LDA model on the cell cycle scores, and calculating cell cycle accuracy. We determined that cell cycle label predictions were 88% accurate (3180 of 3602 cells) (Fig. S6A). Of the 422 cells predicted in an inaccurate cell cycle phase, 419 cells (99.3% of the 422) were predicted to be in an adjacent cell cycle phase (Fig. S6A-B). Given the nature of the continuous cell cycle trajectory being discretized for prediction on cell cycle scores, we speculated that cells predicted in an adjacent cell cycle label may not be incorrect, but instead, likely transitioning. If these cells are removed or counted as correct due to this ambiguity, the model achieves an accuracy of 99.9%. Thus, the cell cycle LDA model is not overfitting and is an accurate computational representation of the cell cycle.

To dissect the circular topology of the cell cycle LD embedding, we tested the hypothesis that the circularity of the model is dependent on the correlations of the input features. We introduced an equal weight of noise into the cell cycle scores to reduce the strength of the correlations (Fig. S6C-D). Equal reduction in the strength of the correlations did not significantly impact the circular embedding, indicating that weak correlations can sufficiently drive a circular trajectory in LDA if the correlative relationships between adjacent class labels are preserved. While balanced disruption of cell cycle scores to produce weak correlations sufficiently retained a circular embedding, the angular pseudotime estimates fell apart when a significant amount of noise was inserted into the data (Fig. S6E). However, introducing noise into three of the cell cycle scores disrupted the circular, donut shape of the cell cycle model (Fig S6F-H). The imbalanced disruption of cell cycle scores indicated that the correlative relationships of the input features and their neighbor classes are important for generating models of circular trajectories in a linear transformation technique.

To identify patterns in gene expression associated with the continuous process of the cell cycle, we computed the angular pseudotime from a computationally-derived startpoint (see methods), and projected the cyclical trajectory into a linear temporal space (Fig. 6F). To verify that the angular pseudotime represented cell cycle patterns, we plotted the cell cycle scores for each phase along the angular pseudotime and found that they increased and decreased according to their expected transitions (Fig. 6G). We then observed the expression of key cell cycle markers over angular pseudotime. We found that the pseudotime expression of CCNB1, CDK1, TPX2, and other cell cycle genes tracked with prior experimentally proven cell cycle transitions (Fig. 6H). For example, CCNB1 transcription increased during S-phase and peaked nearing the G2/M transition (Farshadi et al., 2019). TPX2, a microtubule associated protein responsible for microtubule nucleation that is integral for mitosis, also reached peak expression before entering mitosis (Stewart and Fang, 2005).

To further leverage prior knowledge of experimental conditions with LDA, we utilized the division ID labels of these CFSE+ sorted cells to deconvolve the cell cycle angular pseudotime across the first two divisions (Fig. 6I). We added 0, 360, or 720 to the angular pseudotime value of each cell corresponding to the cell’s respective 0, 1, or 2 division ID, resulting in a continuous axis of cell cycle pseudotime that accounts for the number of divisions a cell has undergone. We observed the gene expression patterning of activated T-cells across these divisions, highlighting the expression of relevant TCR signaling genes such as NFKB and NFATC, as well as cell cycle genes (Fig. 6J). Interestingly, NFATC1 expression increased prior to the first division and began to fluctuate in a cell cycle dependent manner. NFKB1A expression decreased prior to the first division, briefly increased in G1/S of the first division, then began to decrease again. Consistent with our aggregate analysis of cell cycle gene expression patterns, CCNB1 expression was low in non-proliferating cells and increased with TCR stimulation, eventually following the same strict cell cycle dependent pattern in both Division 1 and Division 2 as cells were proliferating. This serves as an illustration that LDA visualization can be used to derive biologically-relevant information beyond initial class labels. This method is robust to noise in cell cycle scores, and computation of cell cycle LD axes and angular pseudotime is computationally efficient. Traditional analysis of scRNAseq data removes cell cycle effects, blinding analysis to cell cycle effects. However, the cell cycle is implicated in mediating important biological functions such as cellular differentiation, plasticity, and inflammatory response (Li and Kirschner, 2014; Good et al., 2019; Daniel et al., 2021-biorxiv). Deconvolving the continuous trajectory of the cell cycle in single-cell RNAseq data empowers researchers to explore highly resolved cell cycle related biological trends. In summary, we found that LDA can visualize cyclical trajectories in scRNAseq data and that experimentally-derived metadata can be leveraged to further deconvolve single-cell patterns across cell cycle and cell division.

## Discussion

Here, we applied supervised dimensionality reduction by LDA to visualize and explore a range of single-cell datasets generated by mass cytometry, RNA-seq, and ATAC-seq. We implemented HSS to identify combinations of features that optimally separate *a priori* classes, providing biologically-interpretable axes suitable for visualization as well as other downstream analyses. We benchmarked the performance of LDA against UMAP, PCA, and PHATE and found it often, but not always, outperformed other algorithms across datasets and cell counts.

While most dimensionality reduction algorithms are unsupervised, HSS-LDA is a supervised method that identifies and highlights differences between designated cell groups. This supervised approach is inappropriate in situations in which labels are unknown, or in which an unbiased view of the data is preferable. Yet, as many single-cell datasets contain one or more known biological labels, HSS-LDA has wide applicability. Even within a single dataset, the same cells can be visualized across multiple HSS-LDA plots, each highlighting the cellular heterogeneity relevant to a distinct biological label. More generally, supervised computational techniques, including but not limited to LDA, can benefit from information related to experimental design and sample associated metadata. Experiments could be devised such that this metadata could specifically be leveraged by supervised methods during analysis to derive new biological insights.

Other researchers have previously demonstrated the utility of feature selection within a supervised classification framework to study sparse signals in genomic datasets, and our Hybrid Subset Selection is one of many feature selection approaches amenable to discriminant analysis (Lê Cao et al., 2011; Witten and Tibshirani, 2011; Clemmensen et al., 2012). However, these reports do not discuss the local or global structure of single-cell visualizations by these supervised tools and do not demonstrate the breadth of visualization applications to single-cell datasets. Here, we have complemented LDA with an interpretable feature selection method (HSS), showed the intuitive visualization patterns of LDA-based visualizations, highlighted the utility of interpretable features in both magnitude and direction, and shown examples of creative analysis approaches that supervised statistical learning tools for multi-class dimensionality reduction can bring to single-cell biology.

As a linear method, LDA is deterministic, reproducing the same plot from the same cells every time. New, unlabeled data can be visualized on existing LD axes using computationally-inexpensive matrix math. LDA does not capture non-linear relationships, which may limit its performance on some datasets. Our findings, however, reinforce the notion that simple, linear models often perform well, even in the presence of non-linear data. While non-linear methods are indispensable in computational biology, linear methods can perform comparably to more complex machine learning approaches, as others have noted (Christodoulou et al., 2019). Furthermore, as LD axes are linear combinations of expression or accessibility data, LDA delivers the added benefit of yielding biologically interpretable single-cell coordinates.

PCA, the most commonly-used linear dimensionality reduction method, was unsuitable for visualization of some single-cell data. The perceived shortcomings of linear dimensionality reduction for visualization may, however, primarily reflect only the specific shortcomings of PCA. Indeed, PCA suffers from the crowding problem, in which cell coordinates often overlap in two-dimensional space. While LDA can also manifest some degree of crowding, the problem is largely avoided as axes are specifically generated to separate cell groups of interest from one another. UMAP’s solution to crowding is to require a minimum distance between any two cells’ coordinates, which is effective in most situations, but was problematic in our imbalanced datasets. While PCA is not always appropriate for two-dimensional visualization, the algorithm remains a pillar of high-dimensional data analysis. PCA is widely applied in preprocessing of single-cell data and principal components often serve as inputs to non-linear dimensionality reduction methods like UMAP (Luecken and Theis, 2019). Likewise, we found that inputting HSS-LDs into UMAP improved performance over either algorithm alone in the challenging task of visualizing a multi-label, single-cell dataset. LD axes may therefore have added value in processing and analysis of single-cell data, outside of only two-dimensional visualization. This is opposed to non-linear dimensionality reduction that is not typically utilized beyond single-cell data embedding.

We recommend HSS-LDA to be employed on any single-cell datasets in which *a priori* labels are known and expected to segregate cells into somewhat homogenous groups. This includes labeling cell types, cell states, stimulation time points, and spatially-distinct cell subsets. LDA is computationally inexpensive and requires no tuning parameters, so it can be deployed with minimal time investment. This is particularly important for researchers with limited access to high-performance computing resources. LDA can also be applied to visualize biological units other than single cells to tease out differences in summary statistics between samples (Jiang et al., 2021; Moore et al., 2021). HSS-LDA might not perform well in situations in which a large degree of heterogeneity exists within a given class. For example, labeling PBMCs according to the timepoint of origin would likely fail to produce a visualization that segregates timepoints, as each PBMC sample would be composed of many cell types. Instead, comparing individual cell types (*e.g*. monocytes) between time points would be more fruitful. Additionally, while no hard limit exists for the maximum number of distinct cell populations that can be visualized by a single HSS-LDA plot, performance will suffer with an increasing number of populations. This maximum threshold will vary by dataset, but at a minimum, three populations are required for a two-dimensional visualization, as LDA generates [# of classes −1] LDs.

We emphasize that HSS-LDA should not replace other dimensionality reduction methods. We encourage researchers to also apply PCA, UMAP, PHATE, and other algorithms to their datasets to benefit from each algorithm’s unique strengths. Given the diversity of technology used and biology explored by single-cell methods, no dimensionality reduction algorithm is suitable for every situation. Furthermore, visualization is only one aspect of single-cell analysis and should always be supplemented with robust, quantitative, high-dimensional analyses. Supervised dimensionality reduction by HSS-LDA uniquely facilitates interpretable visualization, feature selection, and other downstream analysis utility. We envision HSS-LDA as one of many tools that enable computational biologists to visualize and explore single-cell data.

## Methods

### Dataset transformation and preprocessing

We used only previously published datasets and cell annotations provided by the authors. The CyTOF datasets we used are described in the results and paired with graphical representations. For the Morphometry dataset, we used all cells from a single, healthy donor. For the T-cell metabolic regulome dataset, we used healthy CD8 naive T-cells from the same donor and sampled an equal # of cells for each timepoint. For the Chromotyping dataset, we used all cells across all cell lines. The same cells are used as inputs into each algorithm when making any algorithm comparison. CyTOF datasets were transformed using an arcsinh scale of 5 and percentile normalized using a quantile value of 0.999.

The scRNAseq datasets we used are described in the results and paired with graphical representations. scRNAseq count tables for both the Enterocyte Differentiation and T-cell Proliferation Tracing datasets were extracted from their respective publications. Raw count matrices and corresponding metadata was input in Seurat for downstream preprocessing using the NormalizeData, ScaleData, and RunPCA functions. Data was z-scaled prior to PCA transformation and PC feature matrices were inputted into LDA or UMAP. The scATACseq matrix was similarly processed in Seurat using the Signac extension. LSI reduction was performed prior to input into LDA or UMAP.

### Hybrid-Subset-Selection

The HSS-LDA algorithm uses a stepwise feature selection approach, calculating a separation score for each feature subset, and selecting a final set of features that best separates classes for visualization. The calculated separation score for assessing class separation implements commonly used metrics such as Euclidean distance or silhouette score, as well as pixel-based metrics we have introduced such as pixel density or pixel class entropy (PCE) scores. Users can also define their own separation metric function for use with HSS. We describe the major steps of HSS-LDA below:

1. *Initialize the feature set*: Perform LDA for all pairwise combinations of features and evaluate the separation score. Select the pair of features with the best score as the initial feature set.
2. *Perform forward stepwise selection*: Add each feature to the current feature subset, perform LDA, and evaluate the separation score. Add the feature which results in the best separation score.
3. *Perform reverse stepwise selection*: Subtract each feature in the current feature set, perform LDA, and evaluate the score. Remove the feature from the feature set that most improves the score. If no feature removal improves the score, proceed without removing any features.
4. Repeat steps 2-3 until the feature set contains all features.
5. *Compile the best scores across feature set sizes*: Compile scores for all feature sets evaluated in steps 1-4. For each feature set size (2, 3,… k-1, k features), identify the feature set with the best score.
6. *Compute elbow point and select final model*: Calculate the elbow point of scores against feature size in the list compiled in step 5 to select the final feature set. Perform LDA using that feature set to generate the final model.

The *hsslda* R package includes a vignette tilted “hsslda-intro” to guide users on how to use ‘HSS-LDA for dimensionality reduction, feature selection, visualization, and exploratory analysis.

### Dimensionality Reduction Algorithms

We used 4 dimensionality reduction algorithms including LDA, PCA, UMAP, and PHATE plus our Hybrid Subset Selection (HSS)-LDA approach. Software versions, accession links, and parameters used are listed in Supplementary Table #1.

### Runtime analysis

Runtimes were measured using base R Sys.time() function immediately before and after each algorithm function call to fairly evaluate all algorithm runtimes using the exact same input matrices.

### Data subsetting for quantitative benchmarking

All algorithms were evaluated using the same input cells for each dataset. For subsampling, cells are randomly sampled without replacement, and each subsample was drawn three times for replicate analysis of runtimes. However, a minimum of 20 cells are sampled across each class in a label to preserve the presence of rare populations in datasets with severe class imbalances (eg, blast cells in Morphometry or mitotic cells in Chromotyping).

### Metrics for evaluating separability of cell populations and performing feature selection

Euclidean distance is the distance between the means of each class label and was computed using the stats::dist() R function. It calculates the pairwise euclidean distance of all class means and selects the minimum as the score, only rewarding manifolds that separate all labels. Silhouette score was used to determine class label separability using the silhouette coefficient equation (Rousseeuw, 1987) and was computed using the cluster::silhouette() R function. The closer the silhouette score is to 1, the better the cluster separability. The closer the silhouette score is to −1, the worse the cluster separability.

Pixel Class Entropy (PCE) score is a measure of class label distribution in a biaxial grid. Biaxial plots were pixelated into a 100×100 pixel grid, the entropy of classes in each pixel was evaluated, and the PCE score was the average entropy value of each pixel. Here we defined PCE as 1-[(entropy)/log2(# of unique class labels)]. The closer the PCE score was to 1, the greater the class separation. Pixel density uses the same pixel grid approach as PCE and the percentage of each class label in each pixel was evaluated. The lower the average pixel density score, the better the class label separation. PCE score and pixel density was encoded in the hsslda github page.

### Calculating cell cycle pseudotime start-point and angular pseudotime

The cell cycle’s pseudotime start-point is found by selecting the cell with the largest average G1.S and M.G1 cell cycle score. The angular pseudotime is calculated by assigning a pseudotime value between 1-360 to each cell based on its angular location in the LDA’s cell cycle model. The angular pseudotime of all cells are then adjusted based on the location of the cell at the pseudotime start-point.

### UMAP initialization with HSS-LDA

Initialization of UMAP with HSS-linear discriminants can be accomplished in R by inputting the first two HSS-LDs or LDs as a matrix into the ‘init’ variable of the uwot::umap() function.

### Code availability

HSS-LDA is available with installation instructions in R at https://github.com/mamouzgar/hsslda. Figure generation code will be made available upon publication.

### Data availability

All data used will be published and available on public repositories. The BioRxiv for Kimmey et al., 2022 and Baskar et al., 2022 are in process.

## Credit

Conceptualization: DRG, MA, SCB

Methodology: DRG, MA

Software: MA, DRG

Formal analysis: MA

Resources: RB, IA, SCK, AGT, FJH, DRG, SCB

Data curation: MA, DRG

Writing: MA, DRG

Editing: SCB, DRG, MA, RB, FJH

Project supervision: SCB

Funding acquisition: SCB

Funding sources: MA is supported by the Stanford Immunology training grant, T32 AI007290_37. DRG is an awardee of the Bio-X Stanford Interdisciplinary Graduate Fellowship. IA is an awardee of the Weizmann Institute of Science – Israel National Postdoctoral Award Program for Advancing Women in Science. SCK is supported by the NIH/NIGMS Cell and Molecular Biology Training Grant (T32GM007276). RB is supported by the Stanford Cancer Biology Program, and funding from National Science Scholarship (Ph.D.) from Agency for Science, Technology, and Research (A*STAR).

AGT is supported by a Damon Runyon Cancer Research Foundation – DRCRF (DRG-118-16) and Stanford Department of Pathology Seed Grant. This study was supported by an EMBO Long-Term Fellowship ALTF 1141–2017 (to FJH.), the Novartis Foundation for Medical-Biological Research 16C148 (to FJH.) and the Swiss National Science Foundation SNF Early Postdoc Mobility P2ZHP3-171741 (to FJH.). In addition, we received support from National Institutes of Health 1DP2OD022550-01 (to S.C.B.), 1R01AG056287-01 (to S.C.B.), 1R01AG057915-01 (to S.C.B.) and 1U24CA224309-01 (to S.C.B.).

## Supplementary Figures

**Figure S1:**
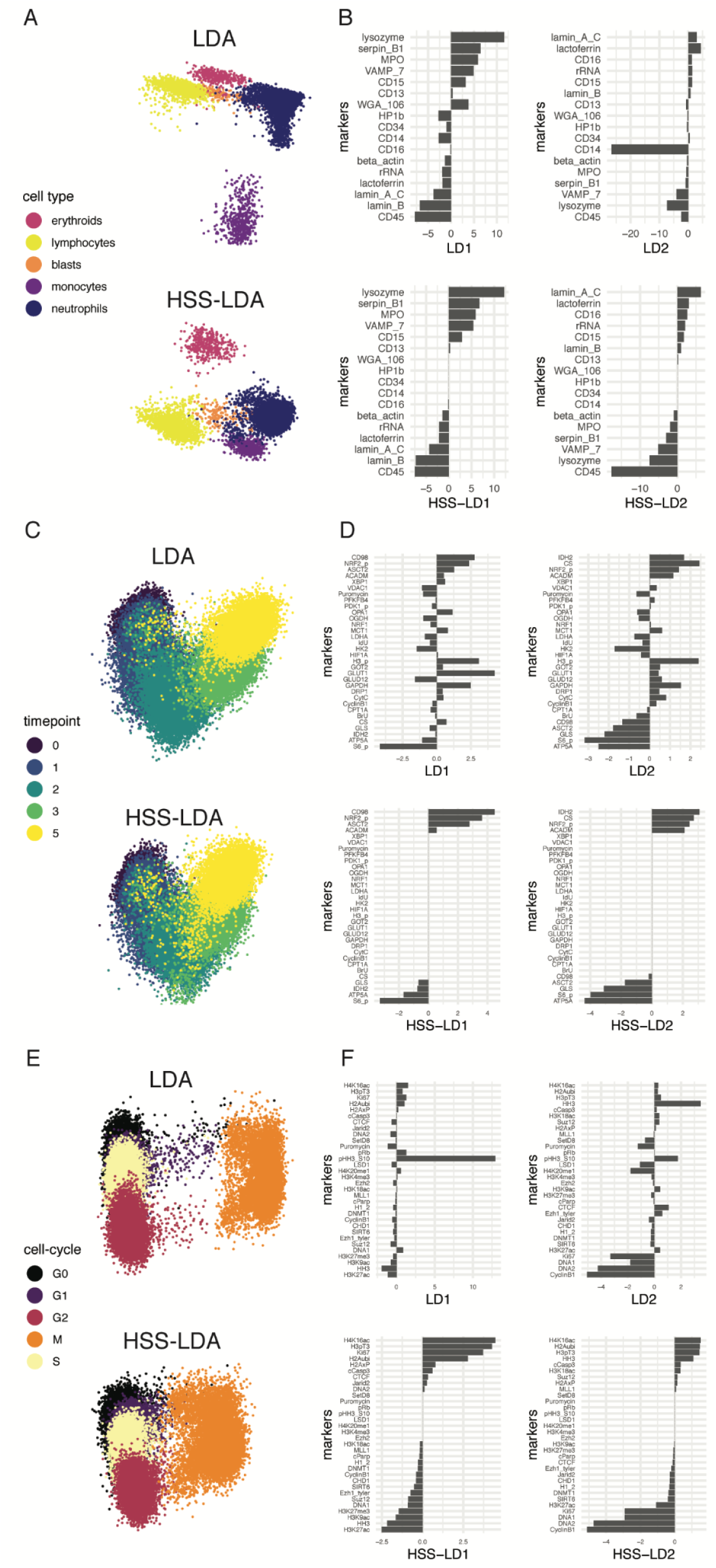
Comparison of LDA to HSS-LDA dimensionality reduction for visualization, feature selection, and feature importance of different mass cytometry datasets. **(A, C, E)** LDA and HSS-LDA embeddings colored by cell-type, collection timepoint, and cell cycle phase for the Morphometry, T-cell metabolic regulome, and Chromotyping datasets, respectively. **(B, D, E)** LDA coefficients and HSS-LDA coefficients for the first two linear discriminants sorted by HSS-LDA coefficient values. Features at zero were removed by HSS-LDA.

**Figure S2:**
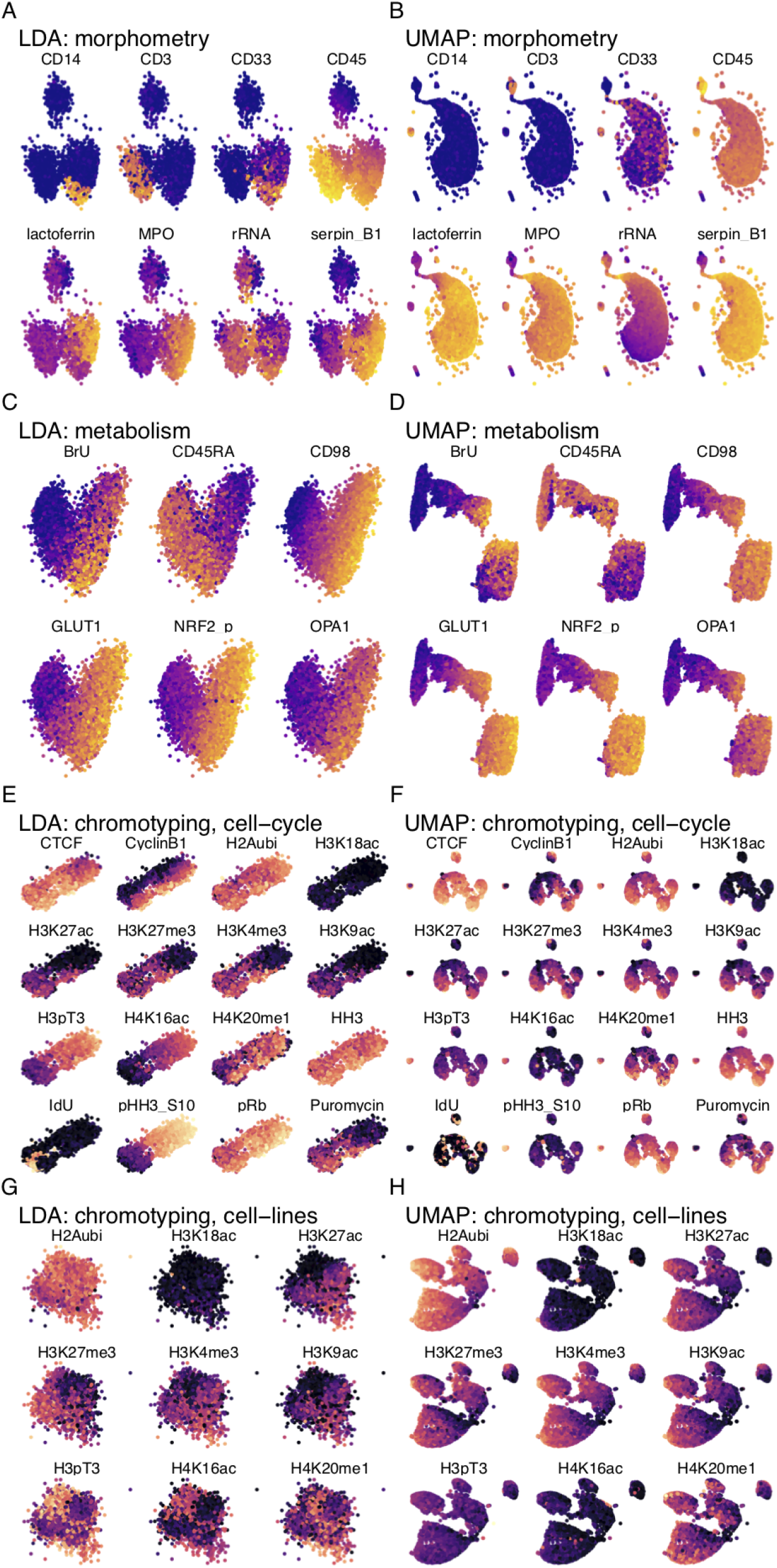
Protein expression of HSS-LDA and UMAP embeddings for different mass cytometry datasets. **(A)** Morphometry HSS-LDA embedding colored by protein expression for cell-type labels. **(B)** Morphometry UMAP embedding colored by protein expression for cell-type labels. **(C)** T-cell metabolic regulome HSS-LDA embedding colored by protein expression for for timepoint labels. **(D)** T-cell metabolic regulome UMAP embedding colored by protein expression for timepoint labels. **(E)** Chromotyping HSS-LDA embedding colored by protein expression for cell cycle labels. **(F)** Chromotyping UMAP embedding colored by protein expression for cell cycle labels. **(G)** Chromotyping HSS-LDA embedding colored by protein expression for cell line labels. **(H)** Chromotyping UMAP embedding colored by protein expression for cell line labels.

**Figure S3:**
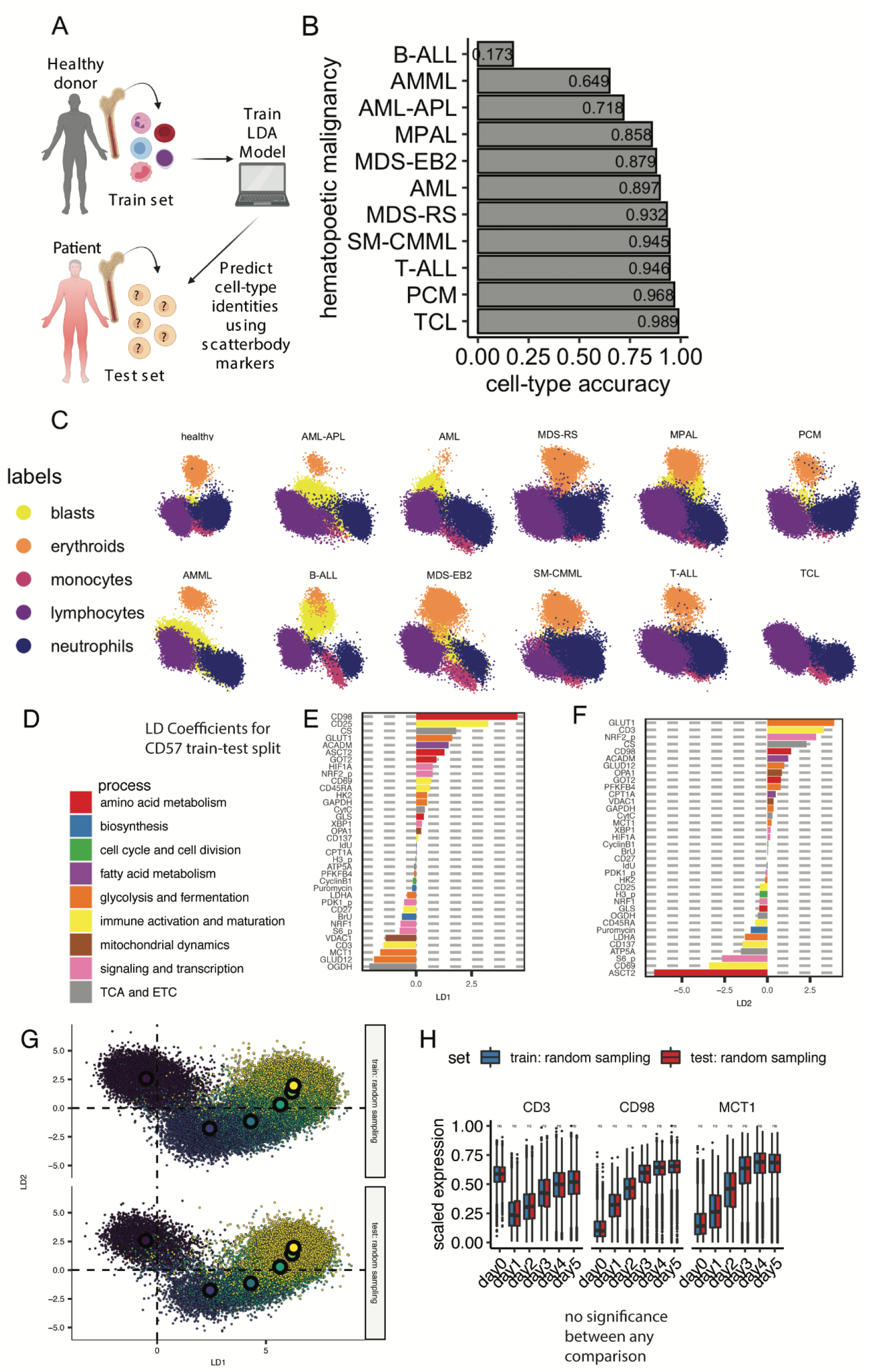
HSS-LDA for predictive analysis of cell-types across different hematopoietic malignancies using scatterbodies, and random sampling of T-cell metabolism dataset for projecting unseen data. **(A)** Graphical illustration of training HSS-LDA model on immune cells from healthy patients using scatterbody protein markers and tested on immune cells from patients with diverse hematopoietic malignancies for accurate cell-type predictions. **(B)** Accuracy of cell-type predictions across different hematopoietic malignancies using HSS-LDA. **(C)** Biaxial LD embeddings of training set (healthy) and test sets (hematopoietic malignancies) projected onto the healthy LD embedding space. **(D)** Labels for different biological system categories of each marker, derived from Hartmann et al, 2020. **(E-F)** Magnitude and direction of coefficients for all markers in the HSS-LD1 (*E*) and HSS-LD2 (*F*) axes, respectively. **(G)** Biaxial HSS-LD plots of randomly sampled cells independent of CD57 expression and labeled with the centroid point for each timepoint. Supervised learning models are known to have decreased performance on test sets. To test the hypothesis that the metabolically slowed trajectory of CD57^high^ cells seen in *G-I* is not due to decreased performance on the CD57^high^ test set, we randomly sample cells for a train-test split and train the HSS-LDA model. The biaxial HSS-LD plots show there is no change between the training set and test set, indicating the metabolically slowed progression phenotype observed in CD57^high^ cells is not dependent on poor test set performance. **(K)** Boxplot summary of protein expression for randomly sampled cells across each timepoint. Wilcoxon signed-rank test performed between randomly sampled cells across each timepoint shows no significant difference in protein expression between the train or test set, indicating the metabolically slowed phenotype in CD57^high^ cells versus CD57^low^ is a true representation of their metabolic trajectory. Wilcoxon signed-rank test performed between CD57^low^ and CD57^high^ cells across each timepoint. *: p <= 0.05; **: p <= 0.01; ***: p <= 0.001; ****: p <= 0.0001.

**Figure S4:**
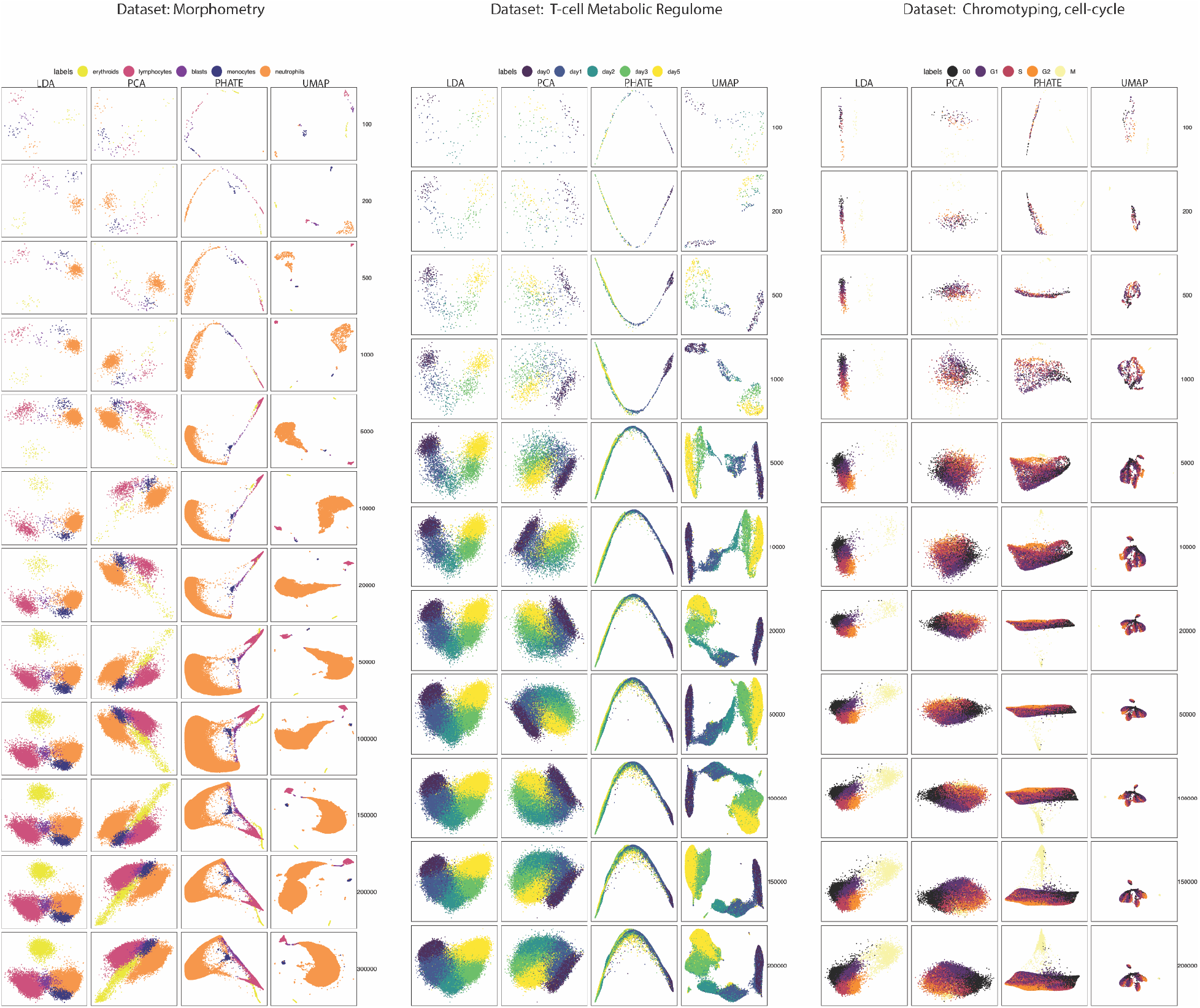
Comparison of biaxial embeddings for LDA, PCA, UMAP, and PHATE across different data subsets. **(A-C)** Biaxial visualizations for varying cell-counts across each algorithm using the **(A)** Morphometry for cell-type, **(B)** T-cell Metabolic Regulome for timepoints, and **(C)** Chromotyping for cell cycle datasets. All algorithms benefit from HSS-LDA feature selection, and the feature matrix for each algorithm is the same for each subset of the data.

**Figure S5:**
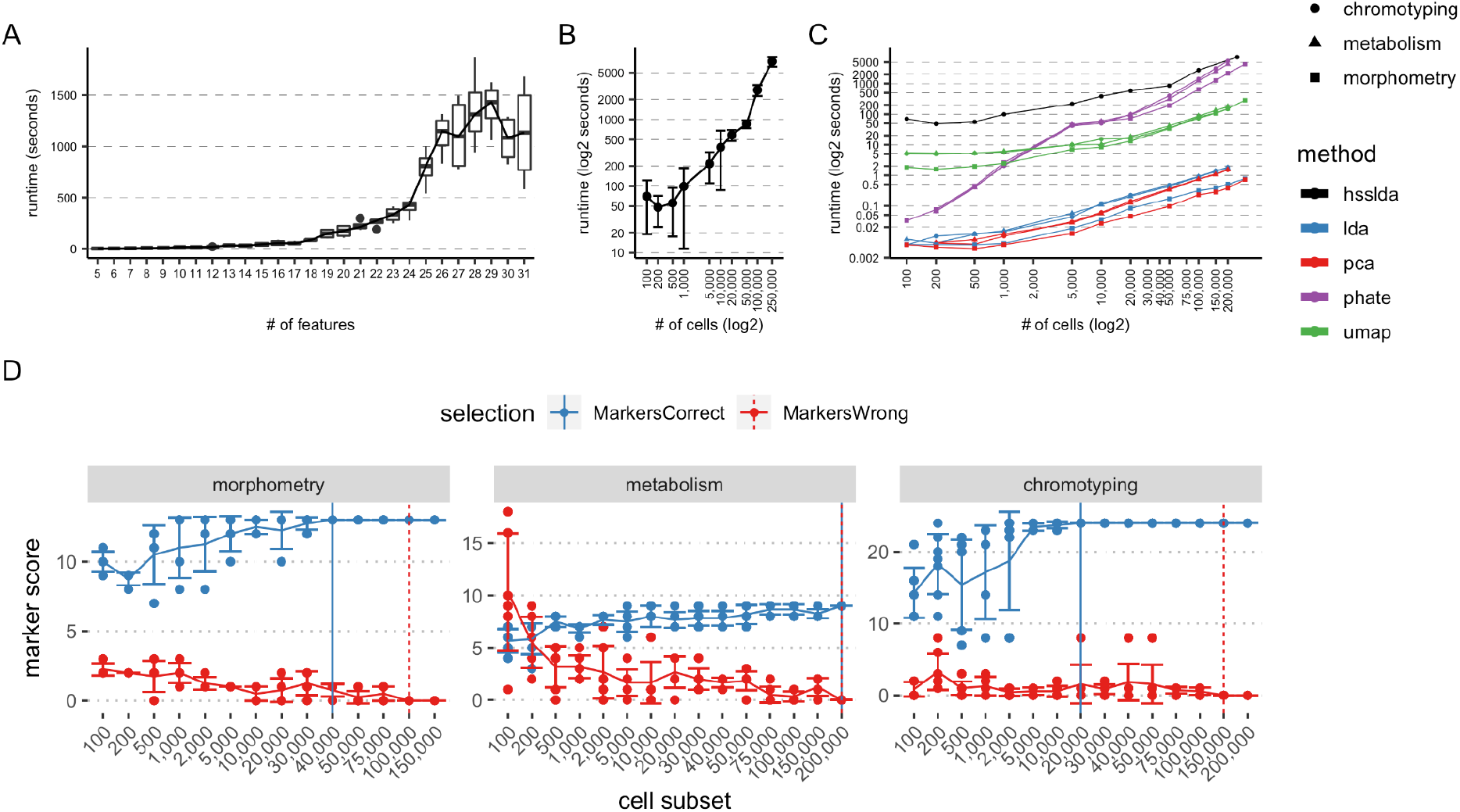
HSS-LDA runtime analysis and assessment of final feature selection. **(A)** Runtime analysis for HSS-LDA with varying # of starting markers randomly selected using the same 50,000 cells. **(B)** Runtime analysis with mean and standard deviation for varying # of cells using 32 chromotyping markers as input feature set. **(C)** Runtime analysis of HSS-LDA benchmarked against all other algorithms that do not perform feature selection. **(D)** Assessment of the minimum # of cells required for HSS-LDA to select the feature set that maximally separates class labels.

**Figure S6:**
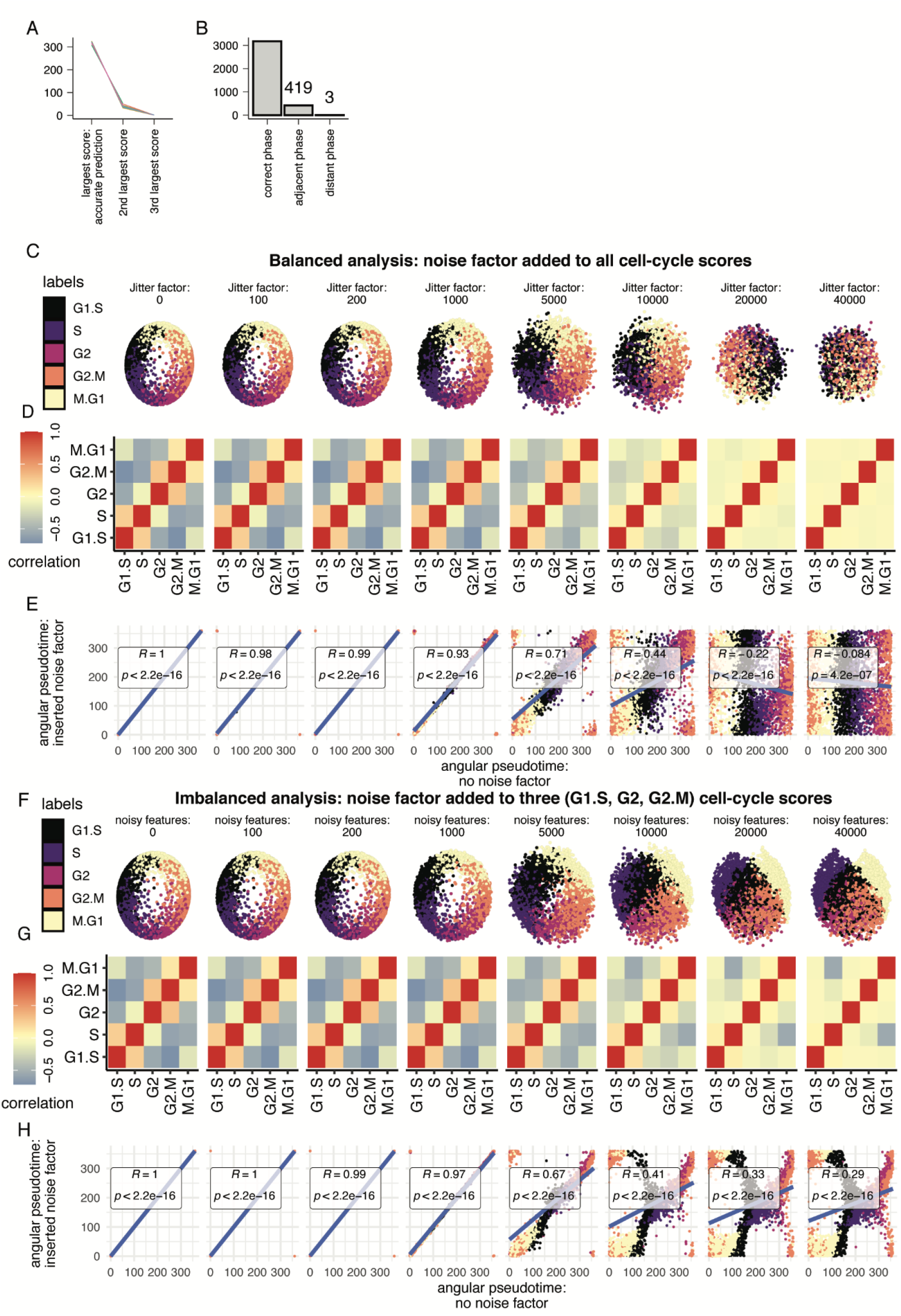
Cross-validation and noise-insertion analysis shows that cell cycle LDA is an accurate embedding of cell cycle progression and that correlative relationships are essential for constructing circular trajectory models in linear transformation techniques. **(A)** Summary barplot of 10-fold cross-validation accuracy results for cyclical LDA model performed on non-overlapping test sets. Accurate cell cycle phase predictions match the largest cell cycle score. 88% test set accuracy. **(B)** Barplot summary of test set predictions indicating cell assignment predictions as either the correct cell cycle phase, an adjacent cell cycle phase, or a distant cell cycle phase. Incorrect predictions are often cells transitioning through the cell cycle in an adjacent cell cycle phase. **(C)** LD embeddings of cell cycle LDAs with increasing amounts of equal noise inserted into all cell cycle scores. **(D)** Correlation heatmaps of cell cycle scores with increasing amounts of equal noise inserted into all cell cycle scores. **(E)** Angular pseudotime estimates of the noisy model versus unperturbed model. A simple linear regression model was generated to determine R2 and P value. **(F)** LD embeddings of cell cycle LDAs with increasing amounts of imbalanced noise inserted into three cell cycle scores (G1.S, G2, G2.M). **(G)** Correlation heatmaps of cell cycle scores with increasing amounts of imbalanced noise inserted into three cell cycle scores. **(H)** Angular pseudotime estimates of the noisy model versus unperturbed model. A simple linear regression model was generated to determine R2 and P value.

## Supplementary Table #1

**Table S1:**
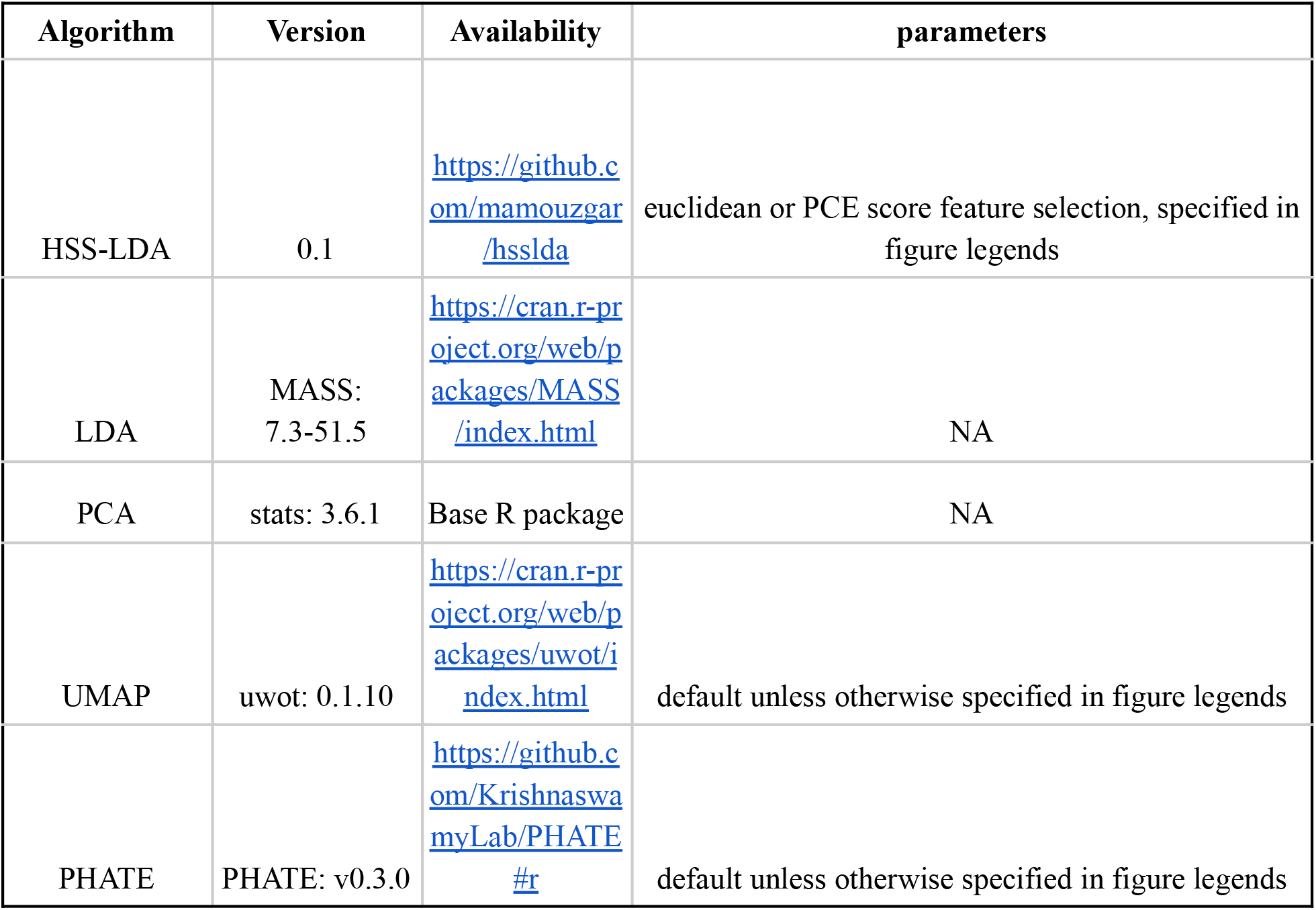
Software versions, accession links, and parameters used for different algorithms

| Algorithm | Version | Availability | parameters |
| --- | --- | --- | --- |
| HSS-LDA | 0.1 | https://github.com/mamouzgar/hsslda | euclidean or PCE score feature selection, specified in figure legends |
| LDA | MASS: 7.3-51.5 | https://cran.r-project.org/web/packages/MASS/index.html | NA |
| PCA | stats: 3.6.1 | Base R package | NA |
| UMAP | uwot: 0.1.10 | https://cran.r-project.org/web/packages/uwot/index.html | default unless otherwise specified in figure legends |
| PHATE | PHATE: v0.3.0 | https://github.com/KrishnaswamyLab/PHATE#r | default unless otherwise specified in figure legends |

